# A *Rhizobiales*-specific unipolar polysaccharide adhesin contributes to *Rhodopseudomonas palustris* biofilm formation across diverse photoheterotrophic conditions

**DOI:** 10.1101/085225

**Authors:** Ryan K. Fritts, Breah LaSarre, Ari M. Stoner, Amanda L. Posto, James B. McKinlay

**Author notes:** Address correspondence to James B. McKinlay,. Present address: Center for Genes, Environment & Health, National Jewish Health, Denver, Colorado, USA.

## Abstract

Bacteria predominantly exist as members of surfaced-attached communities known as biofilms. Many bacterial species initiate biofilms and adhere to each other using cell surface adhesins. This is the case for numerous ecologically diverse *α-proteobacteria,* which use polar exopolysaccharide adhesins for cell-cell adhesion and surface attachment. Here, we show that *Rhodopseudomonas palustris*, a metabolically versatile member of the α-proteobacterial order *Rhizobiales*, encodes a functional unipolar polysaccharide (UPP) biosynthesis gene cluster. Deletion of genes predicted to be critical for UPP biosynthesis and export abolished UPP production. We also found that *R. palustris* uses UPP to mediate biofilm formation across diverse photoheterotrophic growth conditions, wherein light and organic substrates are used to support growth. However, UPP was less important for biofilm formation during photoautotrophy, where light and CO_2_ support growth, and during aerobic respiration with organic compounds. Expanding our analysis beyond *R. palustris*, we examined the phylogenetic distribution and genomic organization of UPP gene clusters among *Rhizobiales* species that inhabit diverse niches. Our analysis suggests that UPP is a conserved ancestral trait of the *Rhizobiales* but that it has been independently lost multiple times during the evolution of this clade, twice coinciding with adaptation to intracellular lifestyles within animal hosts.

**IMPORTANCE:** Bacteria are ubiquitously found as surface-attached communities and cellular aggregates in nature. Here, we address how bacterial adhesion is coordinated in response to diverse environments using two complementary approaches. First, we examined how *Rhodopseudomonas palustris*, one of the most metabolically versatile organisms ever described, varies its adhesion to surfaces in response to different environmental conditions. We identified 2 critical genes for the production of a unipolar polysaccharide (UPP) and showed that UPP is important for adhesion when light and organic substrates are used for growth. Looking beyond *R. palustris*, we performed the most comprehensive survey to date on the conservation of UPP biosynthesis genes among a group of closely related bacteria that occupy diverse niches. Our findings suggest that UPP is important for free-living and plant-associated lifestyles but dispensable for animal pathogens. Additionally, we propose guidelines for classifying the adhesins produced by various *α-proteobacteria*, facilitating future functional and comparative studies.

## INTRODUCTION

Diverse bacteria produce cell surface adhesins that facilitate attachment to biotic and abiotic surfaces (1, 2). Some of the earliest observations of bacterial adhesion reported bacterial ‘stars’, later termed rosettes, in which cells aggregate by attaching to each other at a single pole (3). Similarly, initial observations of bacterial adhesion to abiotic surfaces also noted polar attachment (4). It has since been recognized that the same polar adhesins responsible for rosette formation in many α-proteobacterial species also mediate irreversible attachment to surfaces and thereby act to initiate formation of surface-associated communities known as biofilms (1, 5, 6).

Polar surface attachment in *α-proteobacteria* has been most well studied in the freshwater bacterium, *Caulobacter crescentus* (5, 7–10), and more recently in the plant pathogen, *Agrobacterium tumefaciens* (11–13). The polar adhesin of *C. crescentus* and other members of the order *Caulobacterales* is called holdfast (1, 14). The polar adhesin of *A. tumefaciens* has been termed unipolar polysaccharide (UPP) (11). These two unipolar adhesins are distinct but share certain genetic, biochemical, and functional characteristics (1, 11). Synthesis of both adhesins involves a Wzy-dependent polysaccharide synthesis and export pathway. For holdfast, this pathway is encoded by the holdfast synthesis (*hfs*) gene cluster (8, 15). For UPP, the pathway is partially encoded by the core *upp* biosynthesis gene cluster, with other components encoded separately in the genome (11). The *hsf EFGHCBAD* and *uppABCDEF* gene clusters each have distinct organization and content (i.e., synteny) (Fig. S1). Only *hfsD* and *hfsE* have close sequence similarity to *uppC* and *uppE*, respectively (Table S1), although other genes likely encode functionally analogous proteins between these two gene clusters. A contrasting feature of these two adhesins is that holdfast-mediated adhesion requires proteins encoded by the holdfast anchor (*hfa*) operon, which keeps holdfast attached to the cell (16). No apparent homologs of *hfa* genes are encoded by *A. tumefaciens* (11) or most other *Rhizobiales* species (Dataset S1). Holdfast and UPP also exhibit some biochemical similarity, as both contain *N*-acetylglucosamine (7, 11), allowing the adhesins to be visualized by fluorescence microscopy after staining with the fluorophone-conjugated wheat germ agglutinin (5, 17). Beyond *C. crescentus* and *A. tumefaciens*, polar polysaccharide adhesins are also a common morphological trait across ecologically diverse *α-proteobacteria* (1, 14, 18), especially among *Rhizobiales* species (19–25). However, the genetic and biochemical diversity of the adhesins across this clade is unclear. Furthermore, the potential environment-specific production and/or function of these adhesins remain largely unexplored. Here we examine polar adhesin production by the *Rhizobiales* member, *Rhodopseudomonas palustris*. This purple non-sulfur bacterium was first reported to produce a polar adhesin almost 50 years ago (26), but the genes involved in its biosynthesis were never characterized. Additionally, *R. palustris* is renowned for its metabolic versatility (27), a feature that allowed us to investigate if adhesin production is coordinated with different metabolic modules. We show that the putative *R. palustris uppE* (RPA2750) and *uppC* (RPA4833) orthologs are required for synthesis of a UPP adhesin. UPP is differentially required for *R. palustris* biofilm formation under various conditions, but is particularly influential under photoheterotrophic conditions, in which light energy and organic substrates are used to support growth. Moving beyond *R. palustris*, we also explored whether UPP is associated with different bacterial lifestyles by performing a comparative genomic analysis across diverse *Rhizobiales* species. Our results indicate that UPP is a conserved ancestral trait of the *Rhizobiales*, and that *upp* genes have been independently lost multiple times during the evolution of the *Rhizobiales* clade. Based on our analysis, we propose that genetic synteny of adhesion biosynthesis genes is a valid criterion on which to designate the polar adhesins of various *Rhizobiales* members as ‘UPP’.

## MATERIALS AND METHODS

### Bacterial strains and growth conditions

All *R. palustris* strains were derived from CGA009 (27) and are listed in Table 1. Unless otherwise indicated, *R. palustris* was grown statically in 10 ml of defined photosynthetic medium (PM) (28) in sealed 27-ml anaerobic tubes with argon gas in the headspace. All *R. palustris* cultures were incubated at 30 °C. All phototrophic cultures were illuminated with a 60-W light bulb. For all heterotrophic conditions, PM was supplemented with succinate as the sole carbon source (15 mM in liquid cultures or 10 mM in agar). Incubation in PM with 15 mM succinate and light are henceforth referred to as standard photoheterotrophic conditions. For low phosphate (P_i_) conditions, PM was modified by replacing Na_2_HPO_4_ and KH_2_PO_4_ (12.5 mM each) with equimolar concentrations of Na_2_SO_4_ and K_2_SO_4_. A 1:1 molar mixture of Na_2_HPO_4_ and KH_2_PO_4_ was added for a final P_i_ concentration of 30 μM. For N_2_-fixing conditions, (NH_4_)_2_SO_4_ was omitted from PM and argon was replaced with N_2_. For high salinity conditions, PM was supplemented with 1.5% (w/v) sea salts (Sigma) or NaCl. For chemoheterotrophic conditions, cultures were grown in 10 ml of aerobic PM supplemented with 0.05% yeast extract in addition to 15 mM succinate in 50-ml Erlenmeyer flasks shaken at 225 rpm in darkness. For photoautotrophic conditions, anaerobic PM was supplemented with 60 mM NaHCO_3_ as the inorganic carbon source and 30 mM Na_2_S_2_O_3_ as an inorganic electron donor. Plasmid-harboring *R. palustris* strains were grown with 50 μg/ml gentamicin in liquid culture and 100 μg/ml gentamicin on agar plates. *Escherichia coli* strains used for cloning (DH5-α, S17- 1) were grown aerobically in Luria-Bertani medium supplemented with 15 μg/ml gentamycin when necessary.

**Table 1.** Strains and plasmids used in this study

| Strain or plasmid | Relevant Genotype and/or description | Reference or source |
| --- | --- | --- |
| *R. palustris* |  |  |
| CGA009 | Wild-type strain | (27) |
| CGA4000 | CGA009 derivative; $\Delta uppE$ ($\Delta RPA2750$) mutant | This study |
| CGA4022 | CGA009 derivative; $\Delta uppC$ ($\Delta RPA4833$) mutant | This study |
| *E. coli* |  |  |
| S17-1 | *thi pro hdsR hdsM+ recA*; chromosomal insertion of RP4-2 (Tc::Mu Km::Tn7) | (31) |
| DH5-α | $F^- \lambda^- recA1 \Delta(lacZYA-argF)U169 hsdR17 thi-1 gyrA96 supE44 endA1 relA1 \Phi 80 lacZ \Delta M15$ | Thermo Fisher Scientific |
| Plasmids |  |  |
| pJQ200KS | Gm<sup>R</sup>; *sacB*; *R. palustris* suicide vector; | (30) |
| pJQ-RPA2750 | Gm<sup>R</sup>; *sacB*; Derived from pJQ200KS; deletion vector for *uppE* (RPA2750) | (59) |
| pJQ-RPA4833 | Gm<sup>R</sup>; *sacB*; Derived from pJQ200KS; deletion vector for *uppC* (RPA4833) | This study |
| pGEM | High-copy-number cloning vector for insertion of PCR products | Promega |
| pBBPgdh | Gm<sup>R</sup>; Broad-host-range cloning vector with constitutive *R. palustris gapdh* promoter | (33) |
| pBBP-RPA2750 | Gm<sup>R</sup>; Derived from pBBPgdh; complementation vector for $\Delta uppE$ ($\Delta RPA2750$) | This study |
| pBBP-RPA4833 | Gm<sup>R</sup>; Derived from pBBPgdh; complementation vector for $\Delta uppC$ ($\Delta RPA4833$) | This study |

### *R. palustris* strain construction

All plasmids and primers are listed in Tables 1 and 2, respectively. Deletion vectors for *uppC* (RPA4833) and *uppE* (RPA2750) were generated by PCR-amplification of the genomic regions flanking the gene to be deleted as described (29). PCR product pairs were fused by overlap extension PCR and cloned into pJQ200SK (30). Vectors were introduced into *R. palustris* by conjugation with *E. coli* S17-1 (31) or by electroporation (32). Complementation vectors for *uppC* and *uppE* were generated by PCR–amplification of each gene along with the putative ribosomal binding site. PCR products were cloned into pBBPgdh (33), and complementation and empty pBBPgdh vectors were introduced into *R. palustris* by conjugation with *E. coli* S17-1.

**Table 2.**
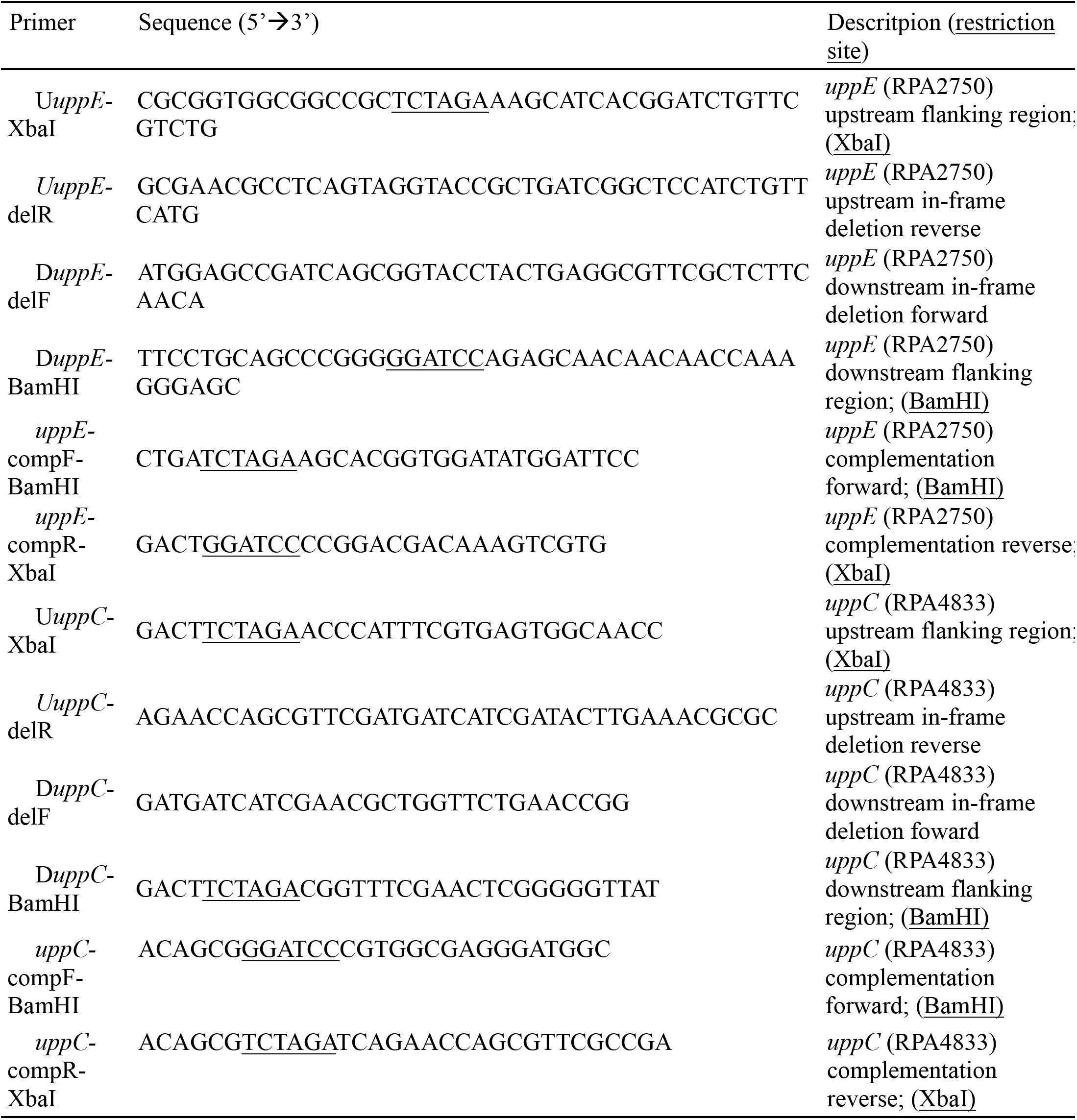
Primers used in this study

### Epifluorescence microscopy and image analysis

Unless stated otherwise, *R. palustris* cultures used for microscopy were grown in liquid without agitation for 2-3 days (d), except for photoautotrophic cultures, which were grown for 8 d. Culture samples were centrifuged and the cell pellet was resuspended in P_i_-buffered saline (PBS) to an optical density between 0.6-0.9 (OD_660_). Wheat germ agglutinin Alexa Fluor^®^ 488 conjugate (WGA-488) (Molecular Probes) was added to cells suspended in PBS at a final concentration of 2 μg/ml and incubated in darkness at room temperature for 15 min. Cells were washed with PBS three times to remove unbound dye and then resuspended in PBS. Cells were imaged on agarose pads using a Nikon Eclipse 90i light microscope equipped with a 100X oil immersion objective and a Photometrics Cascade 1K EMCCD camera, and processed using the Nikon NIS-Elements software. Images were subsequently analyzed using the ImageJ distribution Fiji (34).

### Batch UPP quantification via total WGA-fluorescence

*R. palustris* cultures were grown under standard photoheterotrophic conditions for 3 d to early stationary phase. 400 μl culture samples were centrifuged and the cell pellet was resuspended in 400 μl of PBS. 100 μl of each cell suspension was set aside for use as the unstained control. WGA-488 was added to the remaining 300 μl of resuspended cells to a final concentration of 1.5 μg/ml and incubated in darkness at room temperature for 15 min. WGA488-stained cells were washed three times with PBS and then resuspended in 120 μl PBS to account for cells lost during washes. 100 μl of the stained cells and the reserved unstained samples were each transferred to empty wells of a black polystyrene 96-well μClear^®^ flat bottom microtiter plate (Greiner Bio-One). Fluorescence (top–120: excitation 485/20; emission 528/20) and OD_660_ were measured using a Synergy H1 microplate reader (BioTek). Fluorescence readings were normalized to cell densities (RFU/OD_660_) and background fluorescence was removed by subtracting RFU/OD_660_ values of unstained samples from the WGA-488 stained samples.

### Crystal violet microtiter plate biofilm assay

Biofilm formation was quantified using a modified version a crystal violet microtiter plate assay (35). Briefly, starter cultures were grown under standard photoheterotrophic conditions supplemented with 0.1% yeast extract. 1.5 μl of stationary phase culture was used to inoculate the wells of a lidded, untreated polystyrene 24–well plate (Corning) containing 1.5 ml of the specified sterile medium. All plates were incubated statically at 30 °C. For anaerobic phototrophic growth conditions, plates were incubated in a BD GasPak™ EZ container with two EZ Anaerobe Container System Sachets (BD) and illuminated by two 60-W light bulbs, one on either side of the container. For chemoheterotrophic growth, plates were in air in darkness. For all heterotrophic growth conditions, plates were incubated for 4 d. For photoautotrophic growth conditions (and paired heterotrophic controls), plates were incubated for 10 d. After incubation, plates were shaken at 150 rpm for 3 min on a flat-bed rotary shaker to disrupt loosely attached cells. A 400 μl aliquot of culture was removed for quantifying cell density (OD_660_) for normalization. 400 μl of 0.1% (w/v) crystal violet (CV) was added to each well, and plates were incubated statically at room temperature for 15 min. Wells were then washed 3 times with 2 ml of deionized water to remove unbound CV. 750 ml of 10% (v/v) acetic acid was then added to each well, followed by shaking at 150 rpm for 3 minutes to solubilize bound CV. 150 μl of solubilized CV was transferred to a 96-well plate and absorbance was measured at 570 nm (A_570_). Uninoculated wells containing sterile medium were treated the same way as described above to determine background A_570_, which was subsequently subtracted from all A_570_ measurements.

### Identification of orthologous core *upp* gene clusters and phylogenetic analysis

The putative orthologs of the core UPP biosynthesis genes in *R. palustris* CGA009 (GenBank accession number: BX571963.1) were initially identified by reciprocal best hit analysis using the UppABCDEF proteins of *A. tumefaciens* C58 (GenBank accession number: AE007869.2) as the query sequences for a TBLASTN search against the translated nucleotide database of *R. palustris* CGA009. The best hits in *R. palustris* CGA009 were subsequently used as query sequences for a BLASTX search against the proteome of *A. tumefaciens* C58. All putative *R. palustris* orthologs showed > 50% query cover and an E value < 1˟ 10^-20^ (Table S2). Previous studies noted that the core *uppABCDEF* biosynthesis gene cluster is conserved among several *Rhizobiales* species (20, 23), which was confirmed by using BLASTP with *A. tumefaciens* C58 UppABCDEF proteins as query sequences (Dataset S2). Several additional species that encode complete or near complete *upp* gene clusters were also identified using BLASTP (minimum threshold for homology of query cover > 50%, E value < 1 ˟ 10^-10^; Dataset S2).

For phylogenetic analysis, amino acid sequences for 6 conserved housekeeping proteins, GyrA, GyrB, RpoA, RpoB, FusA, and RecA from 26 α-proteobacterial species were individually aligned using MUSCLE (36) with default settings in MEGA6 (37). Gaps and ambiguous sites were removed from alignments using Gblocks (38), with a minimum block length of 10 positions and gaps allowed at a position for no more than half of the sequences. The final concatenated alignment contained 4,379 amino acid positions (92% of the original positions). Phylogeny was inferred for the concatenated amino acid sequence using the maximum likelihood method based on the Le and Gascuel (LG) 2008 model (39) with 4 discrete gamma categories, which allowed for some sites to be evolutionarily invariable, implemented in MEGA6 (37). The LG+G+I model was selected because it was the best-fitting substitution model based on having the lowest Bayesian information criterion score. Node values indicate branch support from 100 bootstrap replicates. Initial tree(s) for the heuristic search were obtained by applying the Neighbor-Joining method to a matrix of pairwise distances estimated using a Jones-Taylor-Thornton model.

### Statistical Analysis

All statistical analyses were performed using GraphPad Prism version 6.07. Additional information about statistical analyses are in the figure legends and for Fig. 3A, in Table S4.

## RESULTS/DISCUSSION

### Genomic organization of the putative *R. palustris* CGA009 core *upp* gene cluster

*R. palustris* has long been known to form rosettes (26, 40), however the genetic loci responsible for polar adhesin biosynthesis remained uncharacterized. Recently, bioinformatic analysis revealed that *R. palustris* encodes a putative *upp* gene cluster (23). We confirmed that *R. palustris* CGA009 encodes a putative *upp* gene cluster using a TBLASTN reciprocal best hits approach with the *A. tumefaciens* C58 UppABCDEF proteins as query sequences. We identified four adjacent genes in *R. palustris* with close identity to *A. tumefaciens uppABDE* (Fig. 1A, Table S2). Candidate orthologs for both *uppC* (RPA4833) and *uppF* (RPA4581) were outside the putative *R. palustris uppABDE* cluster (RPA2753-2750) (Fig. 1A). As expected based on species relatedness, the synteny of the putative *R. palustris upp* gene cluster is more similar to that of *A. tumefaciens* than to the *C. crescentus hfs* gene cluster (Fig. S1). Also similar to *A. tumefaciens* (11), we did not identify any candidate *hfa* homologs in *R. palustris* (Dataset S1).

**FIG 1.**
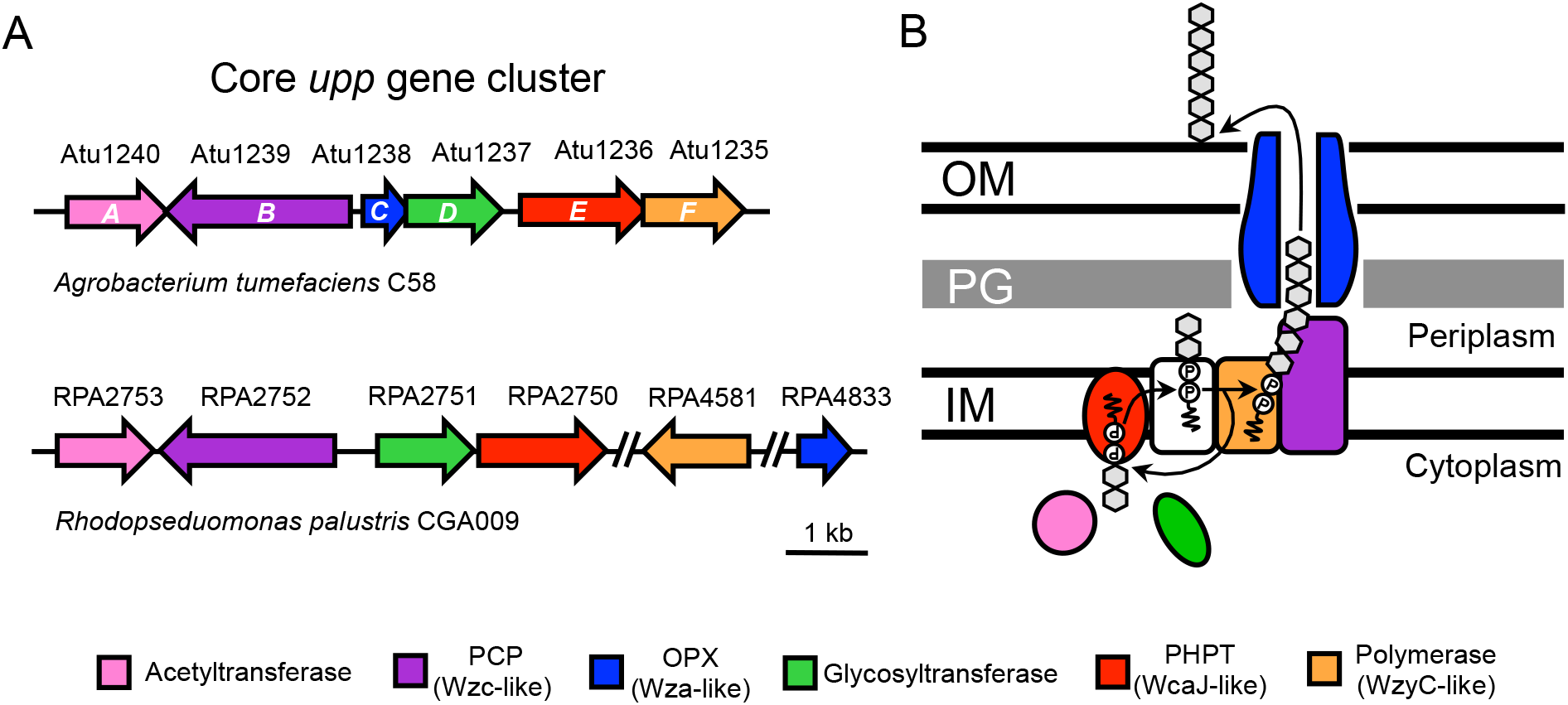
Synteny of *A. tumefaciens* C58 and *R. palustris* CGA009 core *upp* gene clusters and proposed protein functions. (A) Genes (arrows) are colored based on functional prediction and sequence similarity (> 50% query cover, > 25% identity, > 40% positives, and an E value < 1 ˟10^-20^). Double dashes represent large (> 100 kb) unshown genomic regions. (B) Model of the proposed Wzy-dependent synthesis and export pathway for UPP based on (15, 41, 42). *Rhizobiales* core *upp* gene clusters lack an important Wzx-like flippase (white), which is encoded elsewhere in the genome. IM, inner membrane; PG, peptidoglycan; OM, outer membrane; PCP, polysaccharide co-polymerase; OPX, outer membrane polysaccharide export; PHPT, polyisoprenyl-phosphate hexose-1-phosphate transferase.

The putative *R. palustris uppABDE, C, and F* genes are predicted to encode a partial Wzy-dependent polysacchriade export pathway (Fig. 1B). Wzy-dependent pathways are broadly distributed across Gram-negative bacteria (41) and have been most well characterized in lipopolysaccharide and capsular polysaccharide biosynthesis and export in *E. coli* (42). We propose a Wzy-dependent model for UPP synthesis and export based on the current understanding of Wzy-dependent pathways (Fig. 1B), similar to what has been proposed for holdfast production (15). Briefly, an iterative, multi-enzyme process assembles repeat saccharide units (grey hexagons) on the inner membrane (IM)-associated lipid carrier, undecaprenyl phosphate (und-PP). The assembly is then translocated across the IM and into the periplasm where the repeat saccharide units are transferred from und-PP to add to the growing polysaccharide chain on another und-PP carrier. Ultimately the polysaccharide chain is exported onto the cell surface (Fig. 1B). It should be noted that for UPP, certain enzymes thought to be required for synthesis are encoded outside the core *upp* cluster, such as a flippase (Fig. 1B, white) responsible for translocation across the IM. This genetic arrangement is distinct from *C. crescentus* and most other *Caulobacterales* species, which encode putative Wzx-like flippases (HfsF) in their *hfs* gene clusters (15, 17, 43).

### Visualization of *R. palustris* unipolar adhesin

To facilitate genetic and phenotypic characterization of the *R. palustris* adhesin, we first tested if we could visualize adhesin on WT *R. palustris* cells using the fluorophore-conjugated lectin, WGA-488. Adhesins produced by diverse *α-proteobacteria* have been shown to bind WGA (5, 7, 44), which itself binds *N*–acetylglucosamine residues. When we stained *R. palustris* with WGA-488, we observed fluorescence at single poles of some individual cells and at the center of every rosette (Fig. 2A). From this, we conclude that the unipolar adhesin produced by *R. palustris* contains *N*–acetylglucosamine, similar to the UPP of other *Rhizobiales* species (11, 23, 24), as well as *Caulobacterales* holdfast (7, 17).

**FIG 2.**
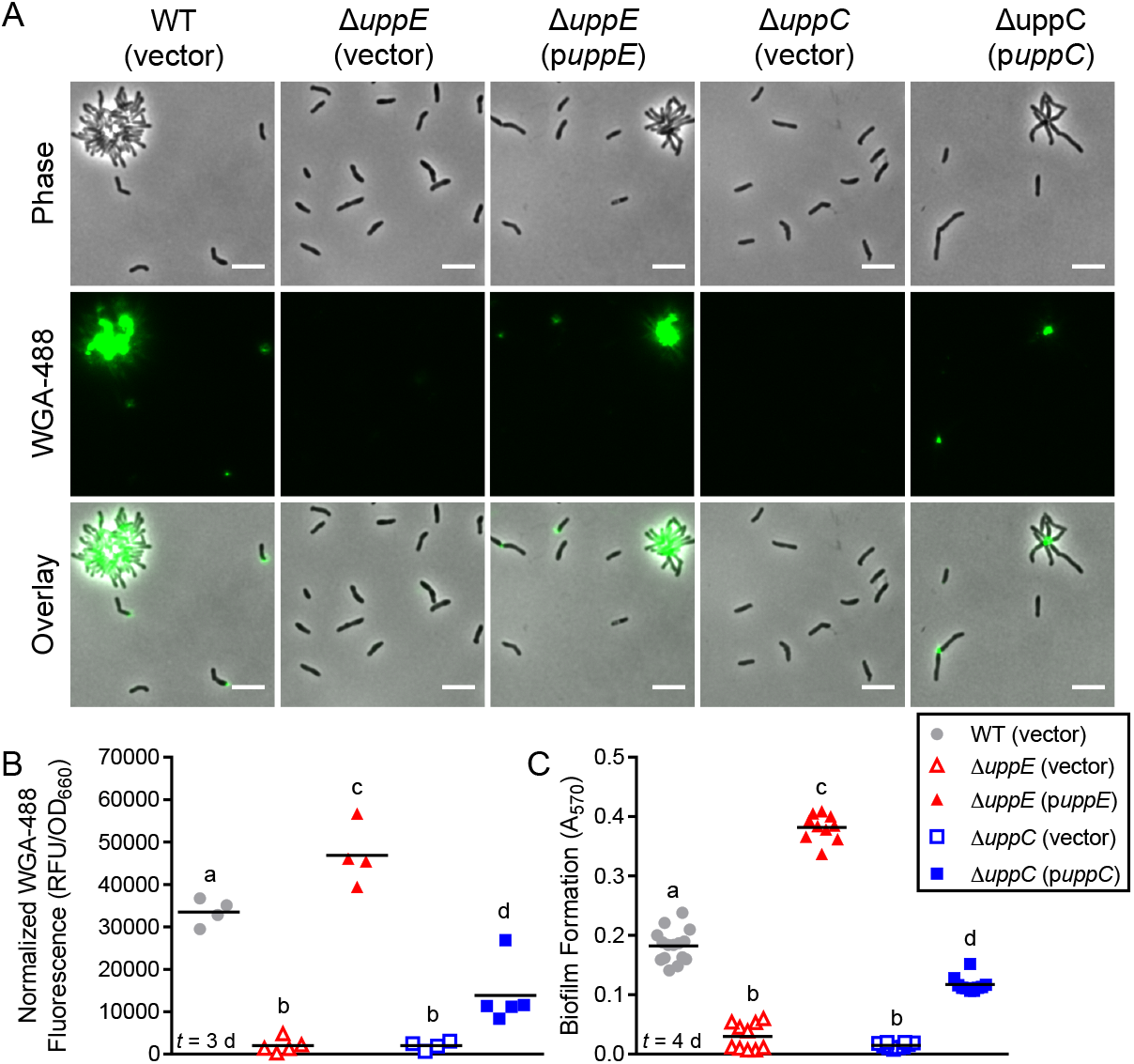
*uppE* and *uppC* are required for UPP biosynthesis, cell-cell adhesion, and biofilm formation. (A) Epifluorescence microscopy of cells stained with WGA-488 after 2 d of growth in standard photoheterotrophic conditions. Scale bars, 5 μm. (B) Normalized total WGA-488 fluorescence from batch UPP quantification following 3 d of growth in standard photoheterotrophic conditions. Different letters indicate significant differences between strains (*P* < 0.05; One-way ANOVA followed by Tukey’s multiple comparisons test; n=4-5). (C) Biofilm formation levels (A_570_) were quantified by CV staining of adherent biomass following 4 d of growth in microtiter wells under standard photoheterotrophic conditions. All strains grew equivalently, so A_570_ values were not normalized. Different letters indicate significant differences between strains (*P* < 0.0001; One-way ANOVA followed by Tukey’s multiple comparisons test; n=10 or 15, pooled from three independent experiments). (B,C) Symbols indicate biological replicates and lines indicate the means. Time (*t*) of sampling following inoculation is indicated in lower left corner.

### UppE and UppC are required for *R. palustris* UPP biosynthesis, cell-cell adhesion, and biofilm formation

We next addressed the genetic requirements underlying polar adhesin production in *R. palustris*. In *A. tumefaciens*, *uppE* (12, 13) and *uppC* (C. Fuqua; personal communication) are essential for UPP biosynthesis. Similarly, the *uppE* ortholog (*gmsA*) of the root-nodulating symbiont *Rhizobium leguminosarum* is necessary for root hair attachment (20). In *C. crescentus,* the putative *uppC* homolog*, hfsD*, is required for holdfast–mediated attachment (8). Thus, we chose the putative *uppE* and *uppC* orthologs of *R. palustris* as targets for in-frame deletions to determine whether they are required for adhesin synthesis.

Deletion of either the putative *uppE* or *uppC* ortholog eliminated both rosette formation and WGA-488 binding (Fig. 2A). Complementation of each mutant from a plasmid restored rosette formation as well as unipolar WGA-488 binding to single cells and at the center of rosettes (Fig. 2A). In addition to microscopic visualization of the adhesin on cells, we also developed an assay to quantify adhesin production at the population level by measuring total WGA-488 fluorescence in batch culture samples. Similar to trends observed by microscopy, total WGA-488 fluorescence was significantly lower in the putative *uppE* or *uppC* mutant cultures compared to WT and the complemented cultures (Fig. 2B). Overall these results demonstrate an essential role for both of these orthologs in adhesin production in *R. palustris*. Based on these results, we henceforth refer to these genes as *uppE* and *uppC*, and to the *R. palustris* unipolar adhesin as UPP.

Having established that *uppE* and *uppC* are critical for *R. palustris* UPP synthesis and251rosette formation, we next assessed if *R. palustris* UPP contributes to biofilm formation. After 4252days of standard photoheterotrophic growth, the *uppE* and *uppC* mutants showed significantly less biofilm formation compared to the WT and complemented strains (Fig. 2C). Thus, we conclude that UPP is the primary adhesin facilitating biofilm formation under standard photoheterotrophic conditions.

### Survey of UPP-mediated biofilm formation across environmental conditions

*R. palustris* is metabolically versatile, allowing it to adopt distinct lifestyles to thrive under diverse conditions. When growing anaerobically in light, *R. palustris* performs anoxygenic photosynthesis to transform energy (27). During phototrophic growth, *R. palustris* can obtain carbon by consuming organic substrates (photoheterotrophy), or by fixing CO_2_, (photoautotrophy) (27). It can also grow by aerobic respiration in the dark (chemoheterotrophy).

Additionally, *R. palustris* is a diazotroph, meaning it can grow with N_2_ gas as the sole nitrogen source, by the process of N_2_-fixation (45). While *R. palustris* has almost exclusively been studied under freshwater conditions, it was recently noted that an environmental isolate could grow in salt concentrations of up to 4.5% (46).

The metabolic versatility of *R. palustris* provided an opportunity to assess whether UPP–mediated surface attachment and biofilm formation is favored by some growth conditions over others. To address this, we examined UPP production and biofilm formation under various growth conditions for both WT *R. palustris* and the *uppE* mutant. We proceeded with only the270*uppE* mutant because we did not observe any phenotypic differences between the *uppE* and *uppC* mutants (Fig. 2). We chose growth conditions that encompass both the metabolic capabilities of *R. palustris* (e.g. N_2_-fixation, photoautotrophy) and abiotic conditions it might normally encounter (e.g. low P_i_, high salinity). Total WGA-488 fluorescence values were not compared across conditions as they were not always reflective of UPP synthesis. For example, some growth conditions, such as low P_i_, resulted in occasional staining at both poles and at what appeared to be cell division septa, suggesting that WGA-488 was staining *N*-acetylglucosomine moieties in peptidoglycan (Fig. S2).

### UPP-assisted biofilm formation is favored by R. palustris in adverse photoheterotrophic environments

We first examined if biofilm formation was stimulated or inhibited in response to three adverse photoheterotrophic conditions. These conditions are considered to be less favorable for *R. palustris* growth due to nutrient limitation (low P_i_), less-preferred nutrients (N_2_-fixation), or osmotic stress (high salinity). Thus, we used these conditions to assess whether biofilm formation might function to increase *R. palustris* survival in suboptimal conditions or to foster persistence in favorable environments (2, 47). We also examined if UPP is utilized by *R. palustris* across these growth conditions. Two main trends were observed under all three adverse conditions. First, WT *R. palustris* formed more biofilm under all adverse conditions compared to standard photoheterotrophic conditions (Fig. 3A), even though standard conditions supported the fastest growth rates and highest cell densities (data not shown). Second, UPP contributed to biofilm formation under all photoheterotrophic conditions, as WT formed more biofilm than the *uppE* mutant in each case (Fig. 3A). These biofilm trends were consistent with microscopy results, which showed that WT *R. palustris* exhibited comparable WGA staining patterns under standard and adverse photoheterotrophic conditions (Fig. 3B). Beyond this, there were also condition-specific phenotypes observed.

**FIG 3.**
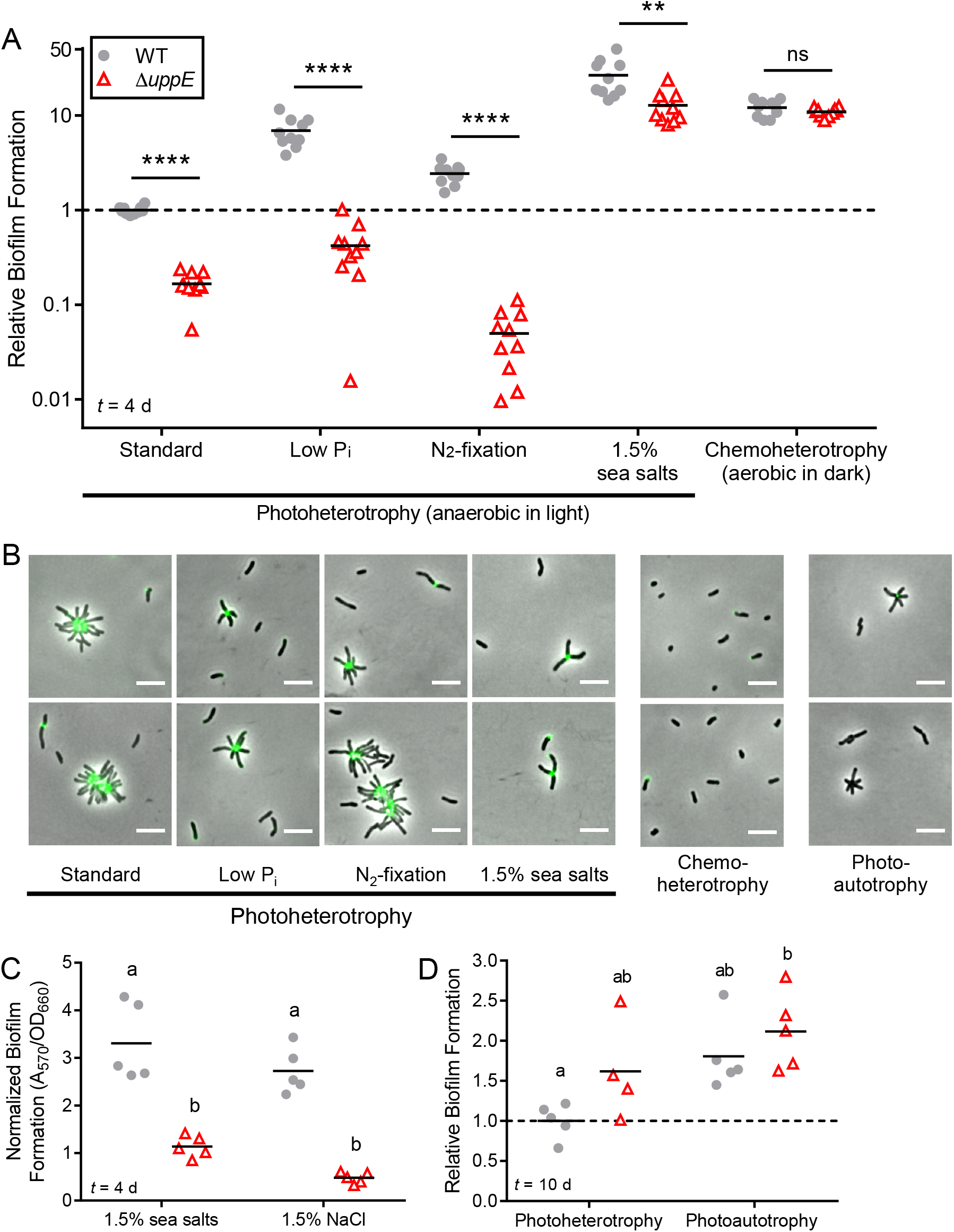
UPP is important for biofilm formation across photoheterotrophic conditions. (A) Biofilm formation levels were normalized to final planktonic cell density (A_570_/OD_660_) and then made relative to normalized WT standard photoheterotrophic values, which was set to 1. \*\**P* < 0.01, \*\*\*\**P* < 0.0001; ns, not significant; based on multiple unpaired, two-tailed t tests without assuming equal variance and followed by Holm-Šídák correction for multiple comparisons; n=10, pooled from two independent experiments. Significance is only indicated for pairwise comparisons between WT and the *uppE* mutant within each condition because the assumption of homogeneity of variances was violated in comparisons across conditions. Results from other statistical analyses comparing across conditions are listed in Table S3. (B) Epifluorescence microscopy of cells stained with WGA-488 after 3 d of photoheterotrophic or chemoheterotrophic growth and after 8 d of photoautotrophic growth. Scale bars, 5 μm. (C) Biofilm formation normalized to final planktonic cell density (A_570_/OD_660_) following 4 d of photoheterotrophic growth with 1.5% sea salts or 1.5% NaCl. Different letters indicate significant differences between groups (*P* < 0.05; Two-way ANOVA followed by Tukey’s multiple comparisons test; n=5). (D) Relative biofilm formation (A_570_/OD_660_) after 10 d of photoheterotrophic or photoautotrophic growth, with WT values from standard photoheterotrophic conditions set to 1. Different letters indicate statistically significant differences between groups (*P* < 0.05; Two-way ANOVA followed by Tukey’s multiple comparisons test; n=4-5). (A, C, D) Symbols indicate biological replicates and lines indicate the means. Time (*t*) of sampling following inoculation is indicated in lower left corner.

Under low P_i_ conditions, the *uppE* mutant formed loosely-attached lawns at the bottom of microtiter wells. These lawns were easily disrupted and washed away. Such lawns were not formed by the *uppE* mutant under standard conditions and were unlike all WT photoheterotrophic biofilms, which were firmly-attached to the sides and bottom of the wells. The genetic and biochemical basis for these loose biofilms remains to be determined. Stimulation of biofilm formation in response to P_i_ limitation has also been observed in *A. tumefaciens* (12, 48). This common observation raises the possibility that increased biofilm formation is a conserved response to P_i_ limitation across some *Rhizobiales* species. It has been speculated that low P_i_ serves as a signal to *A. tumefaciens* that plant surfaces are nearby, as plants sequester P_i_, locally depleting it from the rhizosphere (48). Given that no symbiotic association between *R. palustris* and plants has been identified, it is possible that biofilm formation serves a different function in this species, such as increasing survival when essential nutrients such as P_i_ are limiting.

We also observed 2-fold higher biofilm levels by WT under N_2_-fixing conditions compared to standard conditions (Fig. 3A). N_2_ fixation is energetically expensive compared to using other nitrogen sources such as NH_4_^+^ and is therefore tightly regulated (45, 49). We hypothesize that increased aggregation under N_2_-fixing conditions might function to help retain costly NH_4_^+^, which can passively diffuse out of cells as NH_3_ (50).

In contrast to all other photoheterotrophic conditions, *uppE* mutant biofilm levels were 13-fold higher under 1.5% sea salt conditions than WT cells under standard conditions, despite lacking UPP (Fig. 3A). Similar trends were seen with 1.5% NaCl, confirming that the enhanced biofilm formation of both the WT and the *uppE* mutant was due to high salinity and not another component of the sea salt supplement (Fig. 3C). The high *uppE* mutant biofilm levels under high salinity conditions suggests that additional factors besides UPP contribute to this response. Thus, while UPP-mediated surface attachment contributes to robust biofilm formation by *R. palustris* during photoheterotrophic growth, UPP is less crucial under high salinity conditions.

### UPP-independent biofilm formation is stimulated by non-photoheterotrophic conditions

We also examined UPP production and biofilm formation under chemoheterotrophic and photoautotrophic conditions. Under chemoheterotrophic conditions, UPP was not necessary for biofilm formation, as WT and the *uppE* mutant formed similar levels of biofilm. We were surprised by this result, as it suggested that biofilm formation was entirely UPP-independent. Aerobically-grown bacteria typically adhere near the air-liquid interface (35). However, the adherent biomass of both the WT and the *uppE* aerobic biofilms was at the bottom of the microtiter well, suggesting that *R. palustris* might preferentially form biofilms at microaerobic or anerobic zones. In support of this, the adherent biomass was pigmented, indicating production of bacteriochlorophyll and carotenoids, which is stimulated in response to low O_2_ (51).

Additionally, chemoheterotrophic conditions seem to favor biofilm formation, as WT and *uppE* biofilm levels were approximately 12-fold higher relative to WT under standard photoheterotrophic conditions. (Fig. 3A). Separately, although WGA-488 staining was observed on some single cells, we did not observe any rosettes under chemoheterotrophic conditions (Fig. 3B). It is therefore possible that UPP is produced but is dispensable for chemoheterotrophic biofilm formation.

During photoautotrophy with sodium bicarbonate as the carbon source and thiosulfate as an electron donor, *R. palustris* has a specific growth rate approximately ¼ that of during photoheterotrophic growth (29, 52). Because of the slower growth, we extended photoautotrophic incubations from 4 d to 10 d to allow cultures to reach similar final densities as those observed after 4 d of photoheterotrophic growth. As a control, we also grew parallel photoheterotrophic cultures with equivalent amounts of carbon and electrons for 10 d (Fig. 3D). Under photoautotrophic conditions, WT and the *uppE* mutant showed similar levels of biofilm formation (Fig. 3D), suggesting that biofilm formation was UPP-independent. Similar trends were seen after 10 d of photoheterotrophic growth (Fig. 3D), unlike results from 4 d photoheterotrophic experiments, where the *uppE* mutant formed less biofilm than WT (Fig. 3A). After 8 d of photoautotrophic growth we observed WT rosettes that stained very little or not at all with WGA-488, suggesting that less UPP is produced or that UPP composition is different under these conditions (Fig. 3B). UPP is thought to mediate the initial irreversible surface attachment of cells, so extending the incubation time might have allowed for accessory adhesins or other factors, such as DNA release following cell lysis, to facilitate attachment. Such factors could also contribute to the increased biofilm formation observed across the different conditions tested herein.

Overall, our survey of *R. palustris* biofilm formation across growth conditions can be summarized as follows. UPP mediates biofilm formation under photoheterotrophic conditions, especially those photoheterotrophic conditions that are less favorable to growth. Certain photoheterotrophic conditions, such as high salinity, involve additional factors that are independent of UPP. Finally, chemoheterotrophic and photoautotrophic conditions also stimulate biofilm formation, but in a manner that appears to be entirely independent of UPP.

### Conservation of core *upp* biosynthesis genes across *Rhizobiales* species

Beyond *C. crescentus*, *R. leguminosarum*, *A. tumefaciens*, and now *R. palustris*, the characterization of polar adhesins in other *α-proteobacteria* has been cursory. Historically, all polar adhesins were referred to as holdfast. However, designation of α-proteobacterial adhesins has been complicated by functional differences. For example, the polar glucomannan adhesin of *R. leguminosarum* plays a unique role in root hair attachment but is not required for attachment to abiotic surfaces (19, 20). The *R. leguminosarum* glucomannan biosynthesis gene cluster is orthologous to the *A. tumefaciens uppABCDEF* cluster, which *A. tumefaciens* uses to attach to both biotic and abiotic surfaces (5, 11, 12). Thus, polar *R. leguminosarum* glucomannan and *A. tumefaciens* UPP are homologous adhesins with functional differences. Also contributing to the ambiguity in classifying previously identified *Rhizobiales* polar adhesins is the compositional diversity (1, 12, 20–22). For example, *A. tumefaciens* UPP contains *N*-acetylgalactosamine in addition to *N*–acetylglucosamine (12), the *R. leguminosarum* glucomannan adhesin contains primarily glucose and mannose (19); the *Bradyrhizobium japonicum* polar adhesin contains galactose and lactose (22), and the *Hyphomicrobium* polar adesin likely contains galactose and mannose (21). We therefore propose that *α-proteobacterial* adhesins be classified according to genetic synteny.

Based on the synteny (Fig. 1) and functional requirement of *upp* orthologs for adhesin production (Fig. 2), we conclude that *R. palustris* produces UPP.

With the criterion of genetic synteny in mind, we explored the phylogenetic distribution and genomic organization of the core *uppABCDEF* orthologs across 22 *Rhizobiales* species, representing the lifestyle diversity of this clade (Fig. 4). The topology of this tree is largely consistent with the *α-proteobacteria* phylogeny inferred from a concatenation of 104 protein alignments (53). Our analysis revealed broad conservation of putative *upp* gene clusters, indicating that UPP is an ancestral trait of the *Rhizobiales* clade. Almost all of the *Rhizobiales* plant symbionts, including the plant pathogen, *A. tumefaciens*, the root-nodulating diazotrophs, *R. leguminosarum*, *S. meliloti, Mesorhizobium loti*, and *B. japonicum*, the stem-nodulating photosynthetic diazotroph, *Bradyrhizobium* sp. BTAi, and the leaf epiphyte, *Methylobacterium extorquens,* encode complete or near complete *upp* gene clusters (Fig. 4). The exception to this trend is the root-nodulating diazotroph, *Azorhizobium caulinodans* (54), which does not encode a *upp* cluster (Fig. 4, Dataset S2). We were also unable to identify a *upp* cluster in *Xanthobacter autotrophicus*, a free-living diazotroph closely related to *A. caulinodans* (Fig. 4). This absence suggests that the *upp* cluster was lost before these lineages split. Despite the absence of a *upp* cluster in *A. caulinodans,* it still appear to produce a polar adhesin and forms rosettes (25). Upon closer examination of the *A. caulinodans* ORS571 genome, we identitified a putative Wzy-like polysaccharide biosynthesis gene cluster with high similarity to the *Vibrio fischeri* symbiosis polysaccharide (*syp*) locus (Dataset S3) (55). These putative *syp* homologs seem to have been acquired horizontally and might have been co-opted for polar polysaccharide synthesis in *A. caulinodans*.

**FIG 4.**
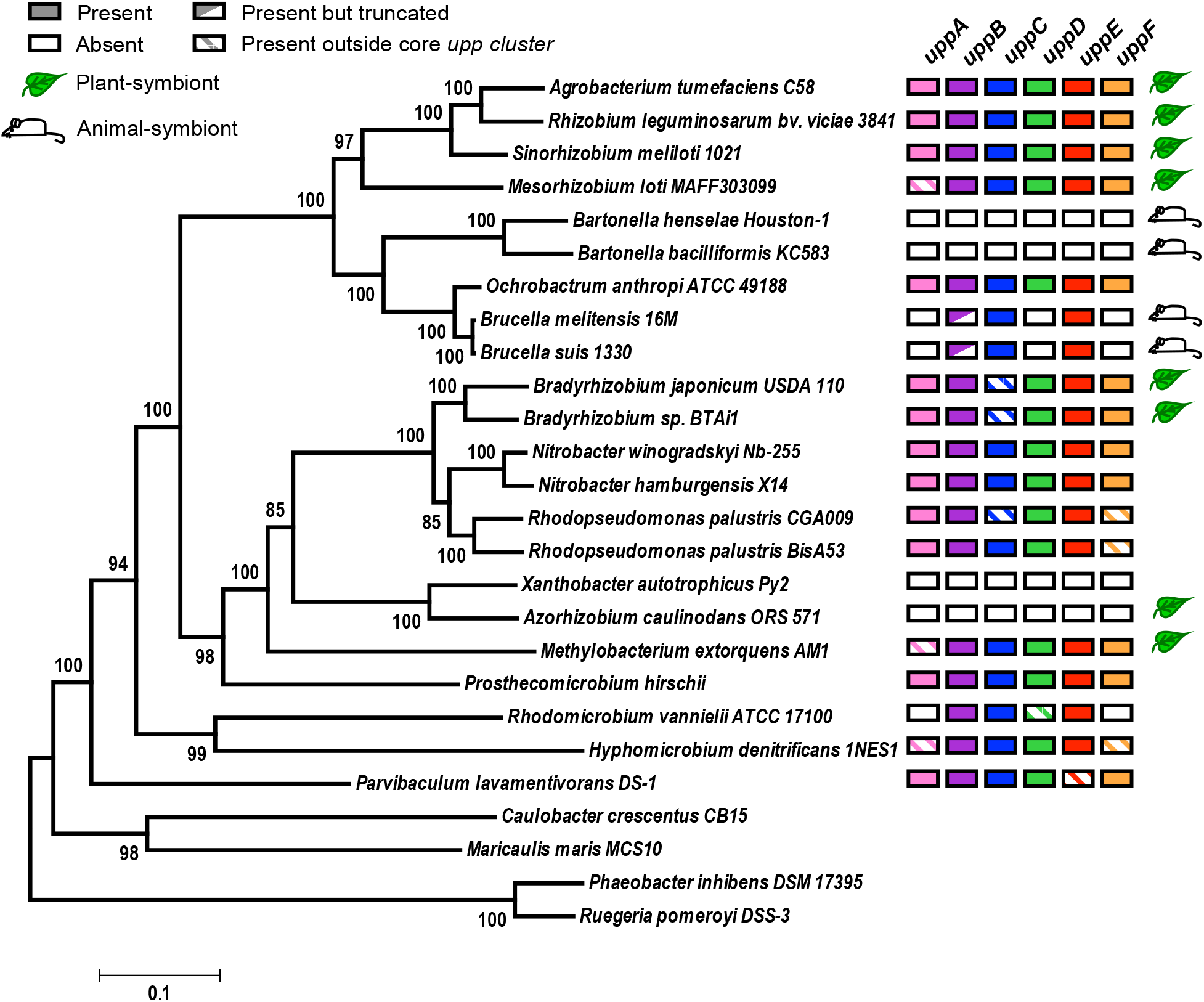
Conservation of core UPP biosynthesis genes among *Rhizobiales* species. A maximum likelihood phylogeny was inferred based on a concatenated alignment of 6 conserved housekeeping proteins using a LG+G+I substitution model (39) with four discrete gamma categories and invariable sites in MEGA6 (37). The tree with the highest log likelihood is shown. Node values indicate branch support from 100 bootstrap replicates. Scale bar represents the number of substitutions per site along branches. Leaf and mouse symbols indicate known plant and animal symbiotic relationships, respectively.

While UPP is well-conserved in plant-associating *Rhizobiales* species, the opposite is true for animal pathogens. This trend was first noted upon the initial discovery of the *upp* gene cluster in *R. leguminosarum,* which noted that this cluster is absent in the *Rhizobiales* intracellular mammalian pathogen, *Brucella melitensis* (20). Rather than being entirely absent (20), our data corroborates more recent bioinformatic evidence that *Brucella* spp. encode a cluster of 3 putative *upp* orthologs (*uppBCE*) (Fig. 4, Fig. S1, Dataset S2) (23). It is not known whether this partial *upp* cluster is involved in the synthesis of a functional UPP. In the closely related intracellular animal pathogens of the genera *Bartonella, upp* orthologs are completely absent (Fig. 4, Dataset S2). In contrast, the soil-dwelling, opportunistic human pathogen *Ochrobactrum anthropi* (56), which is more closely related to *Brucella* than *Bartonella*, encodes a complete *uppABCDEF* gene cluster (Fig. 4). *Ochrobactrum* spp. are thought to be rhizosphere community members but are capable of infecting animal hosts (56, 57). We hypothesize that the entire *upp* cluster was first lost in the *Bartonella* lineage during adaptation to an intracellular lifestyle after diverging from *Brucella/Ochrobactrum*. More recently the *Brucella* lineage has similarly lost multiple *upp* orthologs during its transition to becoming intracellular pathogens. The independent loss of *upp* orthologs in both *Bartonella* and *Brucella* suggests convergent evolution upon adaptation to intracellular niches within animal hosts, supporting the hypothesis that UPP is not important for such lifestyles. Conversely, the conservation of *upp* orthologs in plant-symbionts and free-living species suggests UPP is beneficial in other diverse environments. Considering this, we hypothesize that *Ochrobactrum anthropi* has retained the complete *upp* cluster because it is typically free-living in the soil and thus benefits from producing UPP.

Unipolar adhesins are also used by *α-proteobacteria* outside of the *Rhizobiales.* In the order Caulobacterales, *hfs* and *hfa* gene clusters for holdfast synthesis are well conserved (17, 19 43). Despite the differences in synteny between the *upp* and *hfs* gene clusters (Fig S1), both encode Wzy-dependent pathways for polar polysaccharide synthesis and export and *uppC* and *uppE* show sequence similarity to *hfsD* and *hfsE*, respectively (Table S1). Because of these similarities, we hypothesize that holdfast and UPP share a common evolutionary origin and that the *upp* and *hfs* loci diversified in genomic organization following the divergence of the *Rhizobiales* and *Caulobacterales* clades.

Other α-proteobacterial species of the marine ‘Roseobacter’ clade within the order *Rhodobacterales* also produce polar adhesins and form rosettes but do not encode either *upp* or *hfs/hfa* homologs (18) (Dataset S1 & S2). The polar polysaccharide adhesin of the Roseobacter species *Phaeobacter inhibens* contains *N*-acetylglucosamine based on WGA-binding, indicating that the biochemical composition is at least somewhat similar to UPP and holdfast (44). In this case, polar adhesion synthesis is encoded on a plasmid, since plasmid curing prevented *P*. *inhibens* rosette formation and diminished attachment to abiotic surfaces and algal cells (6). Furthermore, genetic disruption of the plasmid-encoded putative rhamnose operon lowered biofilm formation (58). Plasmids encoding putative rhamonse operons are widely distributed among other Roseobacter species (58), suggesting that polar polysaccharide synthesis and export in this clade is genetically distinct from that of UPP and holdfast. It is not clear whether acquisition of these plasmids led to the loss of gene clusters similar to either *upp* or *hfs* loci.

While polar polysaccharide adhesins are a common morphological trait across ecologically diverse *α-proteobacteria*, there is considerable genetic, compositional, and functional variation, which likely reflects adaptation to different niches. We propose here that genetic synteny of biosynthetic loci is a suitable criterion on which to base classification of polar adhesins. This criterion bypasses uncertainty arising from compositional differences while highlighting the shared underlying biosynthetic pathway. As such, holdfast and UPP are distinct adhesins despite facile similarities. Likewise, the *A. caulinodans* adhesin and the *Roseobacter* rhamnose adhesins should each receive their own designation, as they are genetically distinct from both holdfast and UPP, as well as from each other. Adoption of a unified classification scheme will facilitate both the comparison of adhesins and the exploration of functional differences within and between adhesin types.

## ACKNOWLEDGEMENTS

We thank Yves Brun for use of microscopy facilities and reagents, Clay Fuqua for sharing unpublished data, and members of the Brun and Fuqua labs for thoughtful discussions.

## SUPPLEMENTAL FIGURES AND TABLES

**A *Rhizobiales*-specific unipolar polysaccharide adhesin contributes to *Rhodopseudomonas palustris* biofilm formation across diverse photoheterotrophic conditions** Ryan K. Fritts, Breah LaSarre, Ari M. Stoner, Amanda L. Posto, and James B. McKinlay688Department of Biology, Indiana University, Bloomington, Indiana, USA

**FIG S1.**
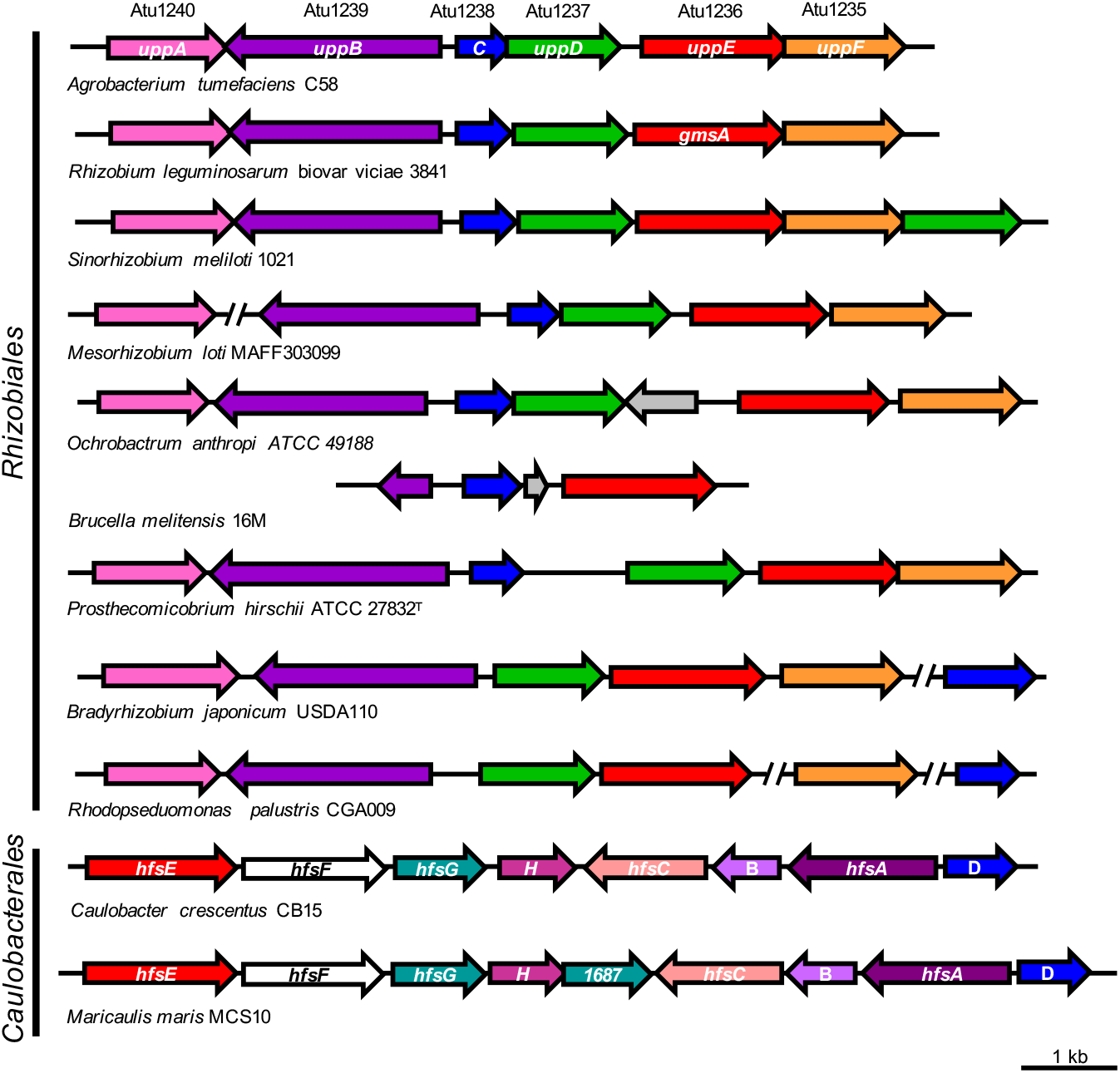
Conservation and synteny of core *upp* and *hfs* gene clusters. Genes (arrows) are colored based on predicted homology and function such that genes of the same color are putative homologs. Genes without a predicted role in polysaccharide biosynthesis and export are shown in gray. Query cover, % Identities/Positives, and E values for and *hfs* and *upp* homologs can be found in Dataset S1 and S2, respectively.

**FIG S2.**
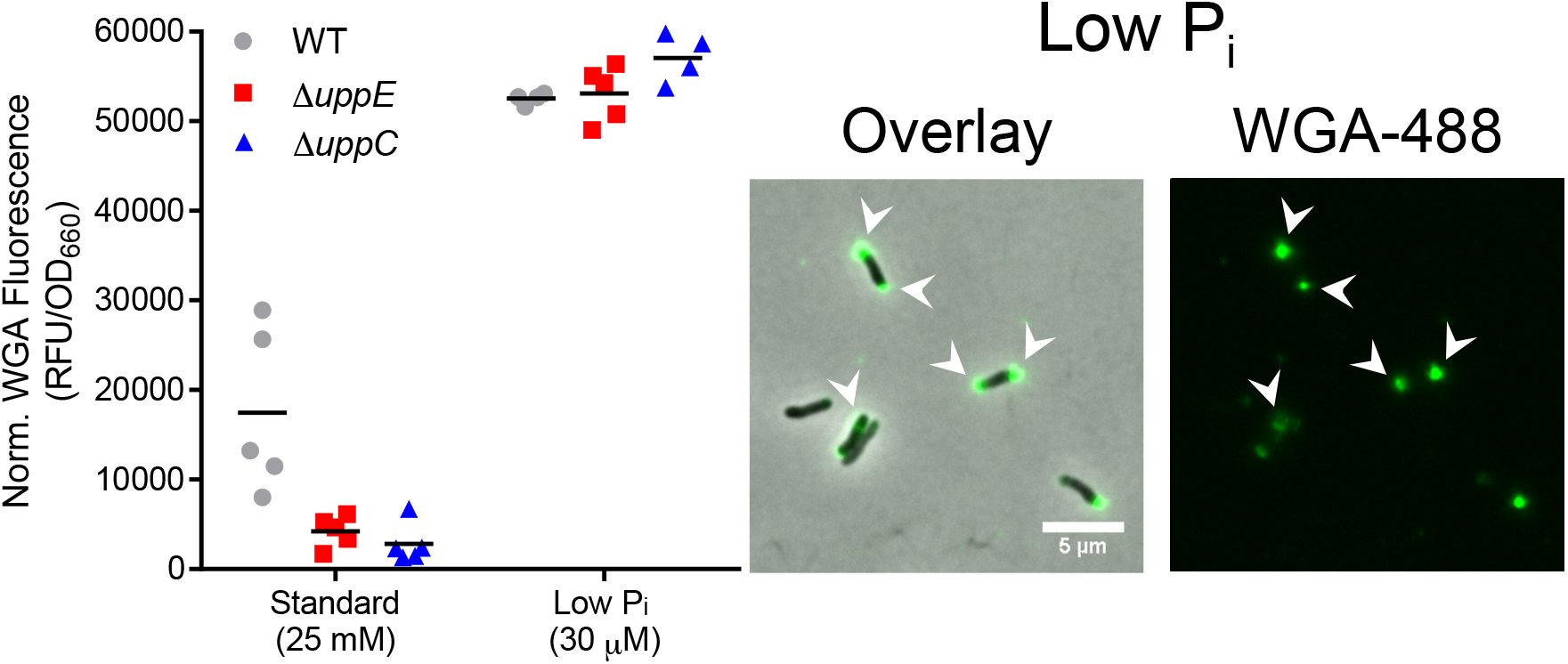
Increased total WGA-488 fluorescence and bipolar and cell body staining of *R. palustris* grown in low P_i_ photoheterotrophic conditions. (A) Normalized total WGA-488 fluorescence from batch UPP quantification via total WGA-fluorescence assay following 3 d of growth under indicated photoheterotrophic conditions. Individual values for biological replicates (n=4-5) are shown with lines indicating the means. (B) Epifluorescence microscopy of WT *R. palustris* cells stained with WGA-488 after 2 days of growth in low P_i_ photoheterotrophic conditions. Arrowheads indicate staining of bipolar or cell body regions. Scale bar, 5 μm.

**TABLE S1.** Putative orthologs of UppABCDEF in *C. crescentus* CB15 identified by reciprocal best hits approach.

| *Agrobacterium tumefaciens*<br>C58 (AE007869.2) query<br>sequence | TBLASTN best hit in *Caulobacter crescentus*<br>CB15 (AE005673.1) |  |  |  | BLASTX reciprocal best hit in *Agrobacterium*<br>*tumefaciens* C58 ((AE007869.2) |  |  |  |
| --- | --- | --- | --- | --- | --- | --- | --- | --- |
|  | locus tag | Query<br>cover | % Ident/<br>Positives | E value | locus tag | Query<br>cover | % Ident/<br>Positives | E value |
| Atu1240/UppA<br>(NP_354252.1, 409 aa) | No matches |  |  |  | N/A |  |  |  |
| Atu1239/UppB<br>(NP_354251.1, 753 aa) | CC_0164 | 51% | 30%/51% | 3.00E-37 | Atu3556 | 99% | 29%/52% | 1.00E-41 |
| Atu1238/UppC<br>(NP_354250.1, 190 aa) | CC_0169 | 86% | 39%/61% | 5.00E-34 | Atu1238/<br>UppC | 100% | 39%/61% | 8.00E-38 |
| Atu1237/UppD<br>(NP_354249.1, 393 aa) | CC_3345 | 46% | 24%/42% | 4.00E-03 | Atu2297 | 92% | 53%/65% | 2.00E-49 |
| Atu1236/UppE<br>(NP_354248.2, 517 aa) | CC_2425/<br>*hfsE* | 70% | 38%/57% | 1.00E-71 | Atu1236/<br>UppE | 89% | 41%/60% | 6.00E-68 |
| Atu1235/UppF<br>(NP_354247.1, 413 aa) | CC_1446 | 11% | 31%/48% | 2.10 | Atu3137 | 57%; | 39%/67% | 8.00E-04 |
| BLASTP Alignment of putative homologs<br>UppC & HfsD (AAK24403.1) |  | 63% | 33%/46% | 4.00E-13 |  |  |  |  |

**TABLE S2.**
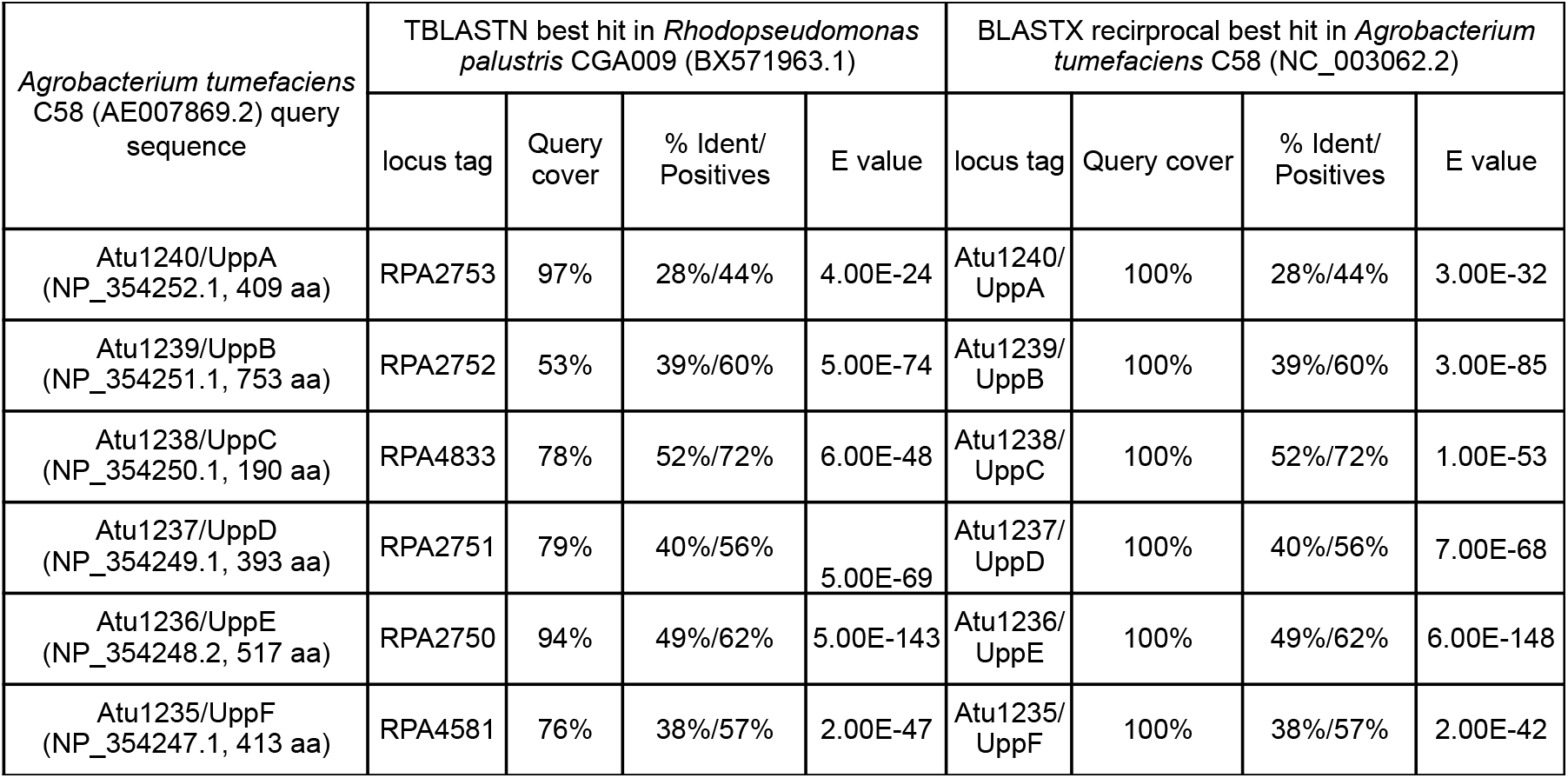
Putative orthologs of UppABCDEF in *R. palustris* CGA009 identified by reciprocal best hits approach.

| *Agrobacterium tumefaciens*<br>C58 (AE007869.2) query<br>sequence | TBLASTN best hit in *Rhodospseudomonas*<br>*palustris* CGA009 (BX571963.1) |  |  |  | BLASTX reciprocal best hit in *Agrobacterium*<br>*tumefaciens* C58 (NC_003062.2) |  |  |  |
| --- | --- | --- | --- | --- | --- | --- | --- | --- |
|  | locus tag | Query<br>cover | % Ident/<br>Positives | E value | locus tag | Query cover | % Ident/<br>Positives | E value |
| Atu1240/UppA<br>(NP_354252.1, 409 aa) | RPA2753 | 97% | 28%/44% | 4.00E-24 | Atu1240/<br>UppA | 100% | 28%/44% | 3.00E-32 |
| Atu1239/UppB<br>(NP_354251.1, 753 aa) | RPA2752 | 53% | 39%/60% | 5.00E-74 | Atu1239/<br>UppB | 100% | 39%/60% | 3.00E-85 |
| Atu1238/UppC<br>(NP_354250.1, 190 aa) | RPA4833 | 78% | 52%/72% | 6.00E-48 | Atu1238/<br>UppC | 100% | 52%/72% | 1.00E-53 |
| Atu1237/UppD<br>(NP_354249.1, 393 aa) | RPA2751 | 79% | 40%/56% | 5.00E-69 | Atu1237/<br>UppD | 100% | 40%/56% | 7.00E-68 |
| Atu1236/UppE<br>(NP_354248.2, 517 aa) | RPA2750 | 94% | 49%/62% | 5.00E-143 | Atu1236/<br>UppE | 100% | 49%/62% | 6.00E-148 |
| Atu1235/UppF<br>(NP_354247.1, 413 aa) | RPA4581 | 76% | 38%/57% | 2.00E-47 | Atu1235/<br>UppF | 100% | 38%/57% | 2.00E-42 |

**TABLE S3.** Statistical analyses for comparing relative biofilm formation by WT across growth conditions in Fig. 3A. Due to violation of the assumption of homogeneity of variances when performing a Two-way ANOVA for the data plotted in Fig. 3A, multiple statistical analyses were performed and compared to reach a consensus for interpreting this data set.

|  | Post-hoc test |  |  |  |  |  |  |
| --- | --- | --- | --- | --- | --- | --- | --- |
| WT only | Dunnett's | Holm-Sidak's | Uncorrected Fisher's LSD | Dunnett's w/o sea salts | Holm-Sidak's w/o sea salts | Dunnett's on log transformed data | unpaired, two-tailed t test (Welch's correction) |
| Standard vs low Pi | 0.0686 | 0.0414 | 0.0209 | < 0.0001 | < 0.0001 | < 0.0001 | < 0.0001 |
| Standard vs N<sub>2</sub>-fixation | 0.9399 | 0.5633 | 0.5633 | 0.1685 | 0.0699 | < 0.0001 | < 0.0001 |
| Standard vs 1.5% sea salts | < 0.0001 | < 0.0001 | < 0.0001 | N/A | N/A | < 0.0001 | < 0.0001 |
| Standard vs chemoheterotrophy | 0.0002 | 0.0001 | < 0.0001 | < 0.0001 | < 0.0001 | < 0.0001 | < 0.0001 |

**TABLE S4.**
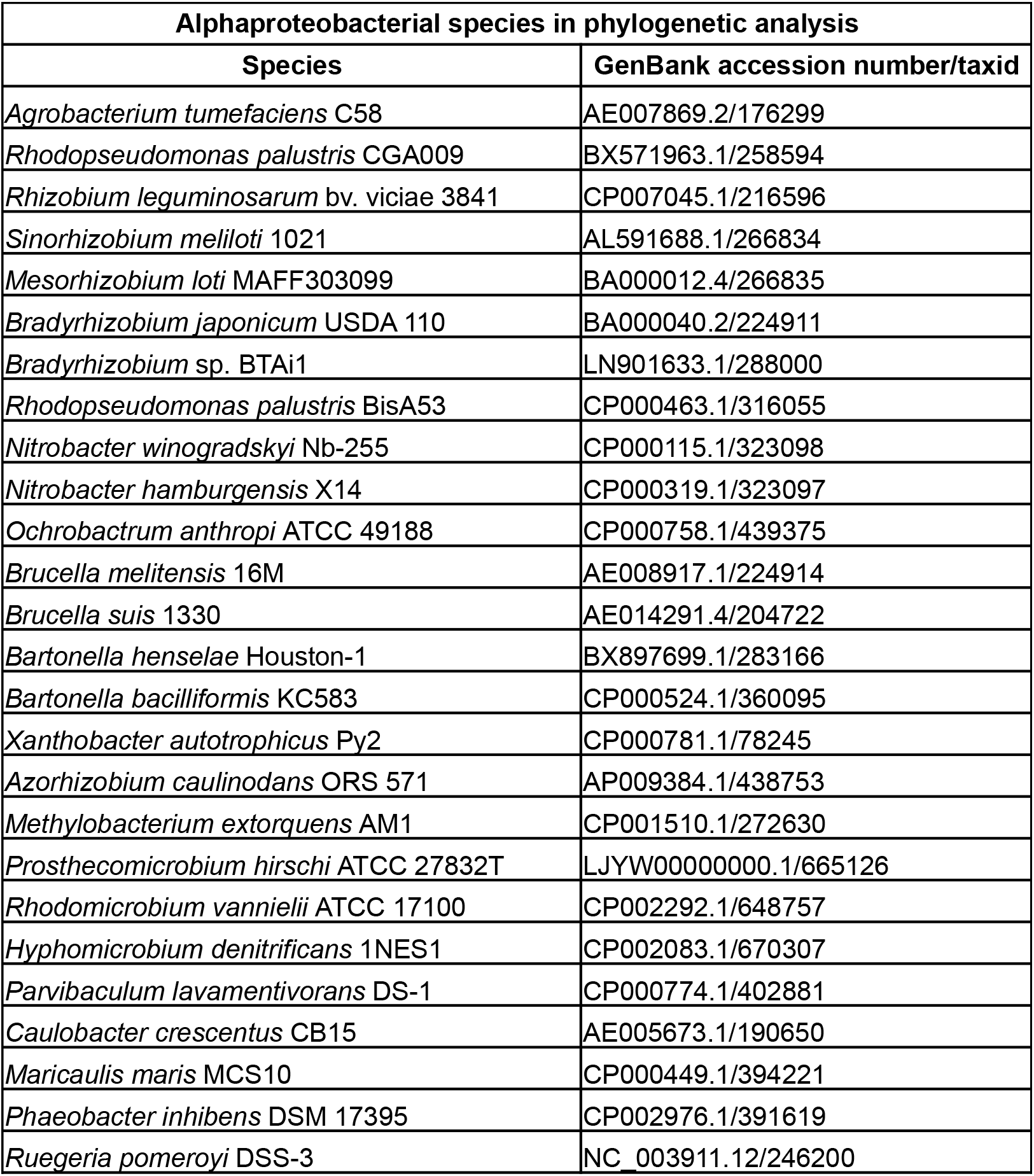
α-proteobacterial species used for phylogenetic analysis. The 26 α-proteobacterial species in the phylogeny in Fig. 4 and their corresponding GenBank accession number and taxid for analysis of the conservation and distribution of core *upp* gene clusters across the order *Rhizobiales.*

